# Gap gene regulatory dynamics evolve along a genotype network

**DOI:** 10.1101/024471

**Authors:** Anton Crombach, Karl R. Wotton, Eva Jiménez-Guri, Johannes Jaeger

**Author notes:** These authors contributed equally to this work (Anton Crombach, Karl R. Wotton). Corresponding authors: Anton Crombach Johannes Jaeger EMBL/CRG Research Unit in Systems Biology Centre for Genomic Regulation (CRG) Dr. Aiguader 88 08003 Barcelona, Spain.

## Abstract

Developmental gene networks implement the dynamic regulatory mechanisms that pattern and shape the organism. Over evolutionary time, the wiring of these networks changes, yet the patterning outcome is often preserved, a phenomenon known as “system drift”. System drift is illustrated by the gap gene network—involved in segmental patterning—in dipteran insects. In the classic model organism *Drosophila melanogaster* and the non-model scuttle fly *Megaselia abdita*, early activation and placement of gap gene expression domains show significant quantitative differences, yet the final patterning output of the system is essentially identical in both species. In this detailed modeling analysis of system drift, we use gene circuits which are fit to quantitative gap gene expression data in *M. abdita* and compare them to an equivalent set of models from *D. melanogaster.* The results of this comparative analysis show precisely how compensatory regulatory mechanisms achieve equivalent final patterns in both species. We discuss the larger implications of the work in terms of “genotype networks” and the ways in which the structure of regulatory networks can influence patterns of evolutionary change (evolvability).

## Introduction

The evolution of biological form involves changes in the gene regulatory networks (GRNs) that underlie organismal development [1–6]. Correspondingly, understanding morphological evolution requires thorough knowledge of the structure of GRNs, the **developmental mechanisms** they encode, and the possible evolutionary transitions between them (terms in bold are defined in Box 1). Over the past sixty years, numerous theoretical and computational studies have led to significant conceptual advances regarding this problem of network evolution; see for example [2–4,7–32]. Yet much remains unknown about network structure and dynamics. On the one hand, subtle alterations in genetic interactions can lead to unexpectedly different regulatory dynamics and hence significant phenotypic changes. On the other hand, major network changes may have no effect on phenotypic output at all. Unfortunately, we do not yet understand the complex and non-linear chain of events that links evolutionary changes in regulatory network structure to changes in developmental mechanisms in any experimentally accessible system [1]. In other words, we know very little—in general terms or in any specific instance—about how the structure of a GRN influences its possible paths of change, its **evolvability** [26,33–35]. Here, we address these issues and supply a first example of a quantitative comparative analysis of developmental GRN structure and dynamics in an experimentally tractable model system: the gap gene network of dipteran insects [36].

##### Box 1: Glossary/Definitions

**Evolvability** can be defined in different ways [35]. In its narrow sense, it describes an evolving system’s propensity for phenotypic innovation; therefore, it has also been called “innovability” in this context [26]. Here, we use it in a more general sense, indicating the capacity of a developmental system to evolve [34]. More specifically, the evolvability of a system reflects the fact that its underlying regulatory network implements a specific set or range of **developmental mechanisms**, and determines the probability of mutational transitions between them. By a developmental mechanism, we mean a collection of regulators and their interactions that generate a reproducible transition from given initial conditions to a specific final state [1,100]. Developmental mechanisms are therefore **dynamic regulatory mechanisms**. They provide causal explanations of how a genotype produces a phenotype (Box Figure A). In this context, **genotype** represents the regulatory structure of network: its components and their interactions; **phenotype** represents the patterning output of the system.

The evolution of developmental mechanisms involves changes in the set of regulators, or changes in their interactions. Such mutational changes can either affect the phenotype of the system or leave it unchanged. **System drift** denotes a mode of network evolution whereby the structure of the network is altered, while the phenotypic output remains constant [5,53–56]. We distinguish between **quantitative system drift**, which affects the strength of regulatory interactions, and **qualitative drift**, which involves recruitment, loss, or exchange of network components as well as rewiring of network structure, either by adding or subtracting interactions, or by changing their signs (activation to repression, or *vice versa*) (see Box Figure B, C, and [46,53,55,56]). It is enabled by the presence of **genotype networks** [26,90], consisting of a set of regulatory network structures that produce the same phenotype, and are connected through small mutational steps (see Box Figure A). In this paper, we examine what kind of regulatory changes produce such a genotype network for the gap gene system in dipteran insects.

**Box Figure.**
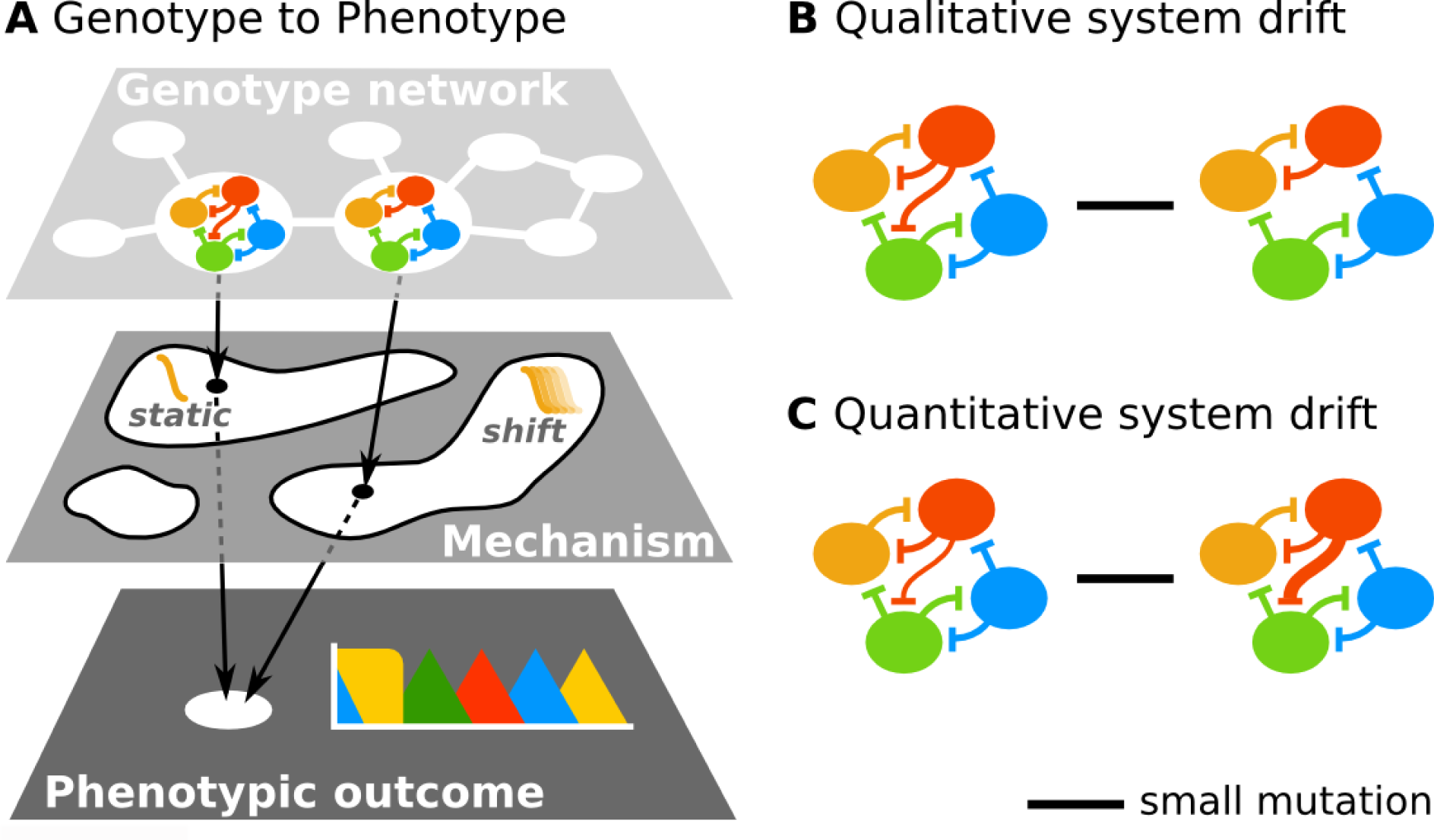

Gap genes are involved in pattern formation and segment determination during the blastoderm stage of early insect development [36]. In dipterans (flies, midges, and mosquitoes, see Figure 1A), they comprise the top-most zygotic layer of the segmentation hierarchy, interpreting maternal gradients to subdivide the embryo into broad overlapping domains of gene expression. We focus on the four key gap genes that operate in the trunk region of the embryo: *hunchback (hb)*, *Krüppel (Kr)*, *knirps (kni)*, and *giant (gt)*.

**Figure 1.**
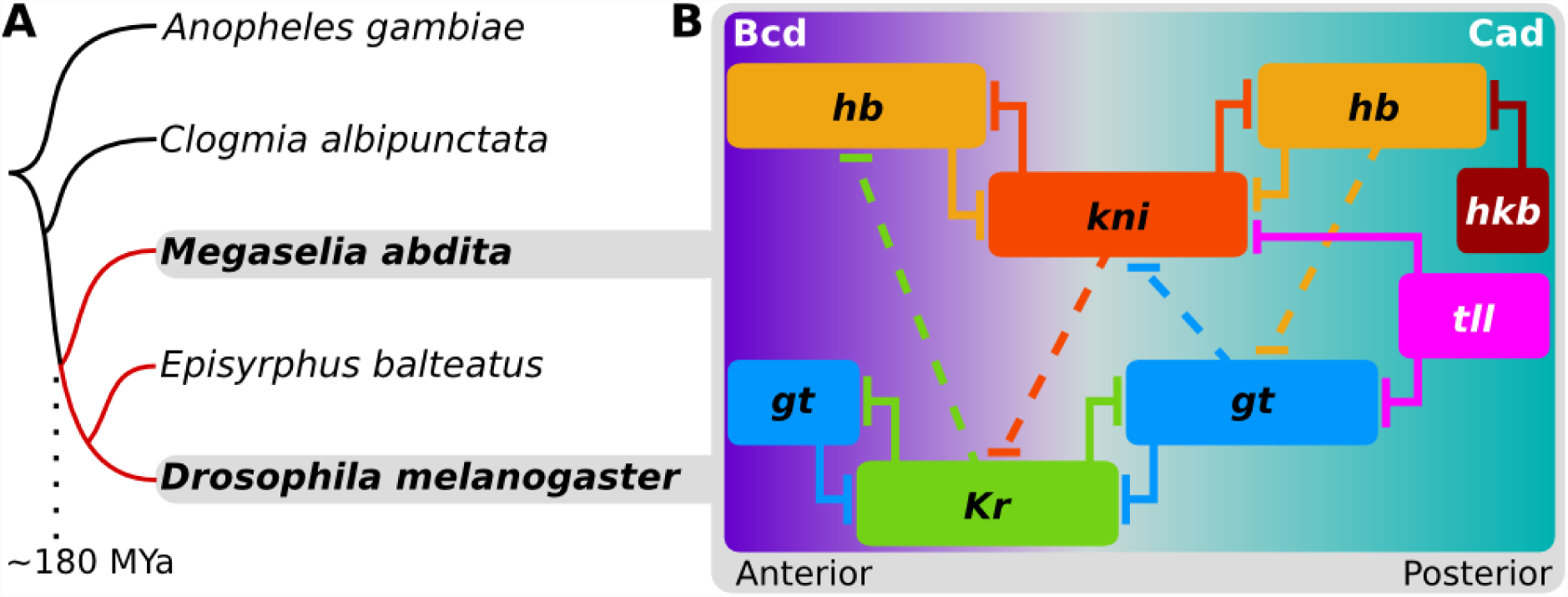
Dipteran phylogeny and structure of the gap gene network. **(A)** Phylogenetic position of *M. abdita* compared to other dipteran species in which gap genes have been studied. Cyclorrhaphan lineage marked in red (MYa: million years ago). **(B)** The gap gene networks of *D. melanogaster* and *M. abdita* share the same qualitative structure. Colored boxes indicate position of gap gene expression domains along the anterior-posterior axis; only the trunk region of the embryo is shown; anterior is to the left, posterior to the right. Trunk gap genes: *hunchback (hb), Krüppel (Kr), giant (gt), knirps (kni)*; terminal gap genes: *tailless (tll), huckebein (hkb)*. Background color represents main activating inputs by maternal morphogen gradients: Bicoid (Bcd) and Caudal (Cad). T-bars represent repression; dashed lines indicate net repressive interactions between overlapping domains.

The developmental mechanisms governing gap gene expression were first worked out in the model organism, *Drosophila melanogaster*. Evidence from genetic, molecular, and data-driven modeling approaches have shown that it implements five basic regulatory principles (Figure 1B) [36]: (i) activation of gap genes by maternal gradients of Bicoid (Bcd) and Caudal (Cad), (ii) gap gene auto-activation, (iii) strong repression between mutually exclusive pairs *hb/kni* and *Kr/gt*, (iv) weak repression with posterior bias between overlapping gap genes causing domain shifts towards the anterior over time, and (v) repression by terminal gap genes *tailless (tll)* and *huckebein (hkb)* in the posterior pole region. In addition to evidence from *D. melanogaster*, gap gene expression and regulation has been studied in a range of non-drosophilid dipteran species [37–49]. This work indicates that the gap gene network is highly conserved within the cyclorrhaphan dipteran lineage of the higher flies (Figure 1B).

In this report, we present a data-driven dynamical modeling approach to analyze and compare regulation of the trunk gap genes between *D. melanogaster* and the non-drosophilid scuttle fly *Megaselia abdita*, a member of the basally branching cyclorrhaphan family Phoridae (Figure 1A, [46,47]). We chose *M. abdita* as our system of study because it is experimentally tractable [50] and features a conserved set of gap genes (and upstream regulators) identical to *D. melanogaster* [46].

Previous work has established the basic qualitative similarities of the gap gene networks in these two organisms [46]. Yet, it was also shown that the precise temporal and spatial dynamics of gene expression differ between them [46]. Specifically, in *M. abdita*, it is thought that a broadened Bcd gradient [39,47,51] and absence of maternal Cad [47,52] lead to gap domains appearing more posteriorly, and retracting from the pole later, than in *D. melanogaster*. Strikingly, however, the system compensates those differences to restore expression boundaries to comparable positions at the onset of gastrulation. And in similar fashion, the embryos of both species have identical patterning when segments appear.

The process leading to such compensatory evolution is called **developmental system drift** [5,53–56]. System drift preserves the outcome of a regulatory process (the “**phenotype**”), while interactions within the network (its “**genotype**”) are altered. Our work shows how such developmental system drift is achieved through regulatory changes in the dipteran gap gene system. We discuss our results within the context of the idea of a **genotype network** [19,26]. Genotype networks consist of related GRNs—connected to each other via small mutations—that all produce the same phenotypic outcome.

They provide a powerful explanatory framework to account for the evolvability of the gap gene network through developmental system drift.

## Results

### Modeling the comparative dynamics of gap gene expression

We previously used gene knock-down by RNA interference (RNAi) to identify conserved and divergent aspects of gap gene network structure between *M. abdita* and *D. melanogaster*. This experimental analysis reveals that the qualitative aspects of the network are highly conserved (Figure 1B); only the strength of specific interactions has changed during evolution [46]. In particular, we identified inter-species differences in sensitivity to RNAi for repressive interactions between overlapping gap genes. These interactions are involved in regulating gap domain shifts in *D. melanogaster* [57]. Based on our evidence, we proposed that the gap gene network is evolving through **quantitative system drift** [46]. This hypothesis provides the starting point for our current investigation into the evolution of expression dynamics governed by gap gene regulation.

At first sight, it may be straightforward and reasonable to assume that the quantitative regulatory changes reported in our previous study [46] can account for the compensatory differences in domain shifts between species. However, genetic analysis using RNAi knock-downs has an important limitation: it remains at the level of correlation, and correlation does not imply causation. For example, an RNAi experiment may reveal an interaction that is particularly sensitive to gene knock-down. But it cannot directly reveal the precise causes and effects of this sensitivity in the context of the dynamic interactions between multiple regulators that constitute a developmental mechanism. Using experimental evidence alone, we cannot establish that the postulated regulatory changes are indeed necessary and sufficient to explain the observed interspecies differences in the dynamics of gap domains.

The aim of our study is to transcend this limitation. We use data-driven mathematical modeling to investigate the complex causal connections between altered network structure and changes in developmental mechanisms that drive the observed differences in expression dynamics between *M. abdita* and *D. melanogaster*. Detailed and accurate models of the gap gene network allow us to simulate and analyze the flow of cause and effect through many simultaneous regulatory interactions. To obtain such models we used a reverse-engineering approach, the gene circuit method; gene circuits are well established and have been successfully applied to the study of gap gene regulation in *D. melanogaster* [57–66]. The approach is based on fitting dynamical network models (“gene circuits”), to quantitative spatio-temporal gap gene expression data from wild-type blastoderm embryos. Importantly, the parameters of a gene circuit not only yield the structure of the network, but also enable detailed analysis of the **dynamic regulatory mechanisms** governing pattern formation by the gap gene system. Since gene circuits do not rely on data derived from genetic perturbations, they yield regulatory evidence which is complementary and independent of that provided by RNAi knock-downs.

### *M. abdita* and *D. melanogaster* gap gene circuits

We created an integrated quantitative data set of gap gene mRNA expression patterns—with high spatial and temporal resolution—for the blastoderm-stage embryo of *M. abdita* (Figure 2A). Our data set is based on previously quantified and characterized positions of gap gene expression boundaries in this species [46]. We used these data to fit gene circuits in order to reverse-engineer the gap gene network of *M. abdita* (Figure 2B). We have previously shown for *D. melanogaster* that both mRNA and protein expression data yield gene circuits with equivalent regulatory mechanisms [66], and that post-transcriptional regulation is not necessary for gap boundary positioning [67]. As a reference for comparison, we used published gap gene mRNA expression data [66] to obtain a set of equivalent gene circuits for *D. melanogaster.* For each species, we selected 20 fitting solutions that capture expression dynamics correctly (Figure 2C–E). See Materials and Methods for details on data processing, model fitting, and analysis.

**Figure 2.**
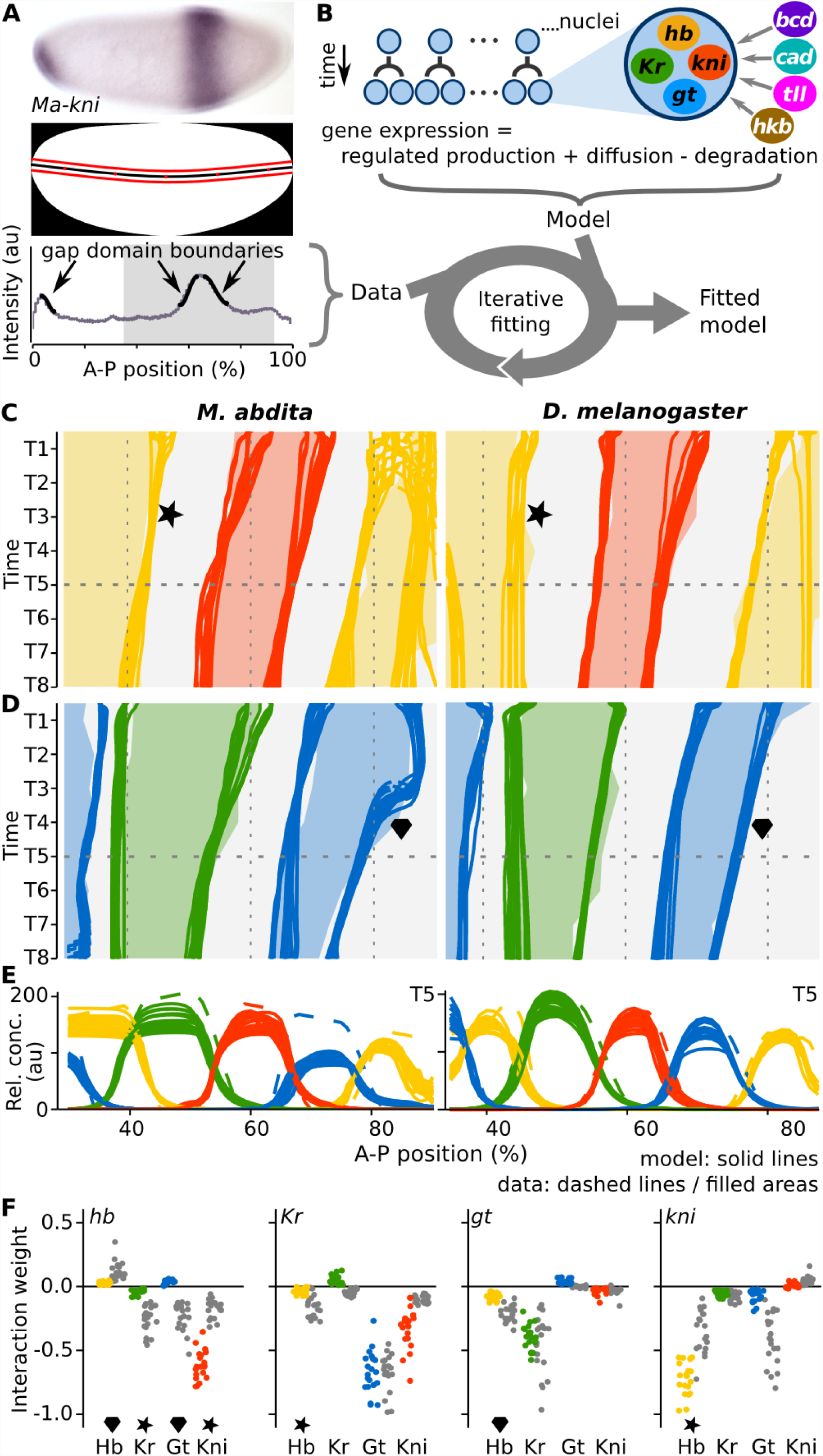
The gene circuit approach and resulting gap gene network models. **(A)** Data acquisition/processing. Top: *M. abdita* whole-mount *in situ* hybridization showing *kni* mRNA expression at mid-blastoderm (C14-T3). Middle: embryo mask showing dorso-ventral midline (black) and 10%-strip used for extraction of expression profiles (bounded by red lines). In both panels, anterior is to the left, dorsal is up. Bottom: extracted *kni* expression profile (grey) in arbitrary units (au); manually fitted spline curves used to extract boundary positions shown in black (arrows); grey background indicates the trunk region included in our models. **(B)** The gene circuit approach: a dynamical model—consisting of a row of dividing nuclei with gap gene regulation, diffusion, and decay—is fit to integrated expression data using a global optimization strategy. **(C–E)** mRNA expression data and gene circuit model output for *M. abdita* (left) and *D. melanogaster* (right) during blastoderm cycle 14A (C14A; time classes T1–8); we show 20 selected gene circuits for each species. **(C, D)** Space-time plots show gap gene expression data (solid areas), overlaid with gene circuit model output (each independent model fit represented by a separate line). Areas/lines demarcate regions with relative mRNA concentrations above half-maximum value. Star indicates dynamic vs. stationary behavior of the posterior *hb* boundary; diamond highlights differing shift dynamics of the posterior *gt* domain. **(E)** Gene expression data (dashed) and gene circuit model output (solid lines) at time class T5 (horizontal dashed line in **C** and **D**). A–P position in percent, where 0% is the anterior pole. **(F)** Comparison of interaction strengths for gap gene cross-regulation between species. Scatter plots show distributions of estimated parameter values from fitted and selected circuits in *M. abdita* (colored dots), and *D. melanogaster* (grey); target genes separated by panel where columns represent regulators. Stars/diamonds indicate interactions involved in corresponding features of expression dynamics highlighted in **C** and **D**.

The resulting models accurately reproduce the observed differences in domain shifts between *M. abdita* and *D. melanogaster* (Figure 2C,D) [46]. This enables us to study the mechanistic basis for these differences through a comparative analysis of gap gene circuits from each species.

### Quantitative changes in conserved gap gene network structure

Gene circuits encode network structure in an interconnectivity matrix of regulatory parameters (see Materials and Methods). We obtain the qualitative structure of the network by classifying the estimated parameter values into categories (activation, no interaction, repression) (Supplementary Figures S2, S3). Our analysis confirms that this qualitative network structure is conserved between *M. abdita* and *D. melanogaster* (Figure 1B), which is fully consistent with evidence from RNAi knock-down experiments [46]. Model analysis reveals that the five basic regulatory principles governing gap gene expression are also conserved: *M. abdita* gene circuits show gap gene activation by maternal Bcd and zygotic Cad, auto-activation, strong repression between *hb/kni* and *Kr/gt*, weaker repression with posterior bias between overlapping gap genes, and repression by terminal gap genes, as in equivalent models for *D. melanogaster* (Supplementary Table S4).

Examining the distribution of estimated parameter values more closely, however, we observe marked quantitative differences in interaction strength between the two species (Figure 2F, Supplementary Figure S6). Many of the altered interactions affect repression between overlapping gap genes, which governs domain shifts in *D. melanogaster* [36,57,66]. Intriguingly, our models predict that these regulatory interactions are often weaker in *M. abdita*, a result which stands in apparent contradiction to previous experimental work [46]. Gene circuit analysis allows us to identify and characterize the precise causal effects of these quantitative changes in interaction strength on the dynamics of gene expression in the complex regulatory context of the whole gap gene network. This enables us to resolve the apparent contradictions between evidence from modeling versus genetic approaches.

### Dynamic *hb* boundary positioned by ratchet-like mechanism

The most salient change in expression dynamics between *M. abdita* and *D. melanogaster* involves the posterior boundary of the anterior *hb* domain (Figure 2C). In *D. melanogaster*, this boundary remains static around 45% A–P position, a fact which is considered crucial for the robustness of gap gene patterning [68–70]. In *M. abdita*, on the other hand, it shifts from 52 to 41% A–P position over time [46].

Regulatory analysis reveals that this qualitative change in dynamical behavior is caused by a combination of altered initial placement of expression domains and changes in relative repression strength. Compared to *D. melanogaster*, the early anterior *hb* domain extends further posterior in *M. abdita*—due to the broader distribution of maternal Bcd [39,46,51]. This results in considerable initial domain overlap between *hb* and *Kr*. The co-existence of these factors across several nuclei is made possible by the limited strength of mutual repression between the two genes (Figure 2F). In return, though, the increased extent of expression overlap leads to a stronger overall repressive effect, since the regulatory contribution of an interaction not only depends on its strength, but also on the concentration of the regulator within the region of overlapping gene expression. It is this overall repressive effect—not the regulatory strength of the interaction as represented by interconnectivity parameters—that is shown in regulatory plots in Figures 3–5.

These plots reveal that within the extended zone of overlap in *M. abdita*, weak repression by *Kr* down-regulates *hb*, whose gradual disappearance eventually allows *kni* to become expressed (Figure 3A,C,E and Supplementary Figure S8; see also Supplementary Text S1). Kni in turn strongly inhibits *hb* (Figure 2F). This results in a ratchet-like mechanism: initial repression by *Kr* primes successive nuclei in the region of the boundary shift to switch irreversibly from *hb* to *kni* expression (mechanisms are summarized in Figure 7). In contrast, much stronger mutual repression between *hb* and *Kr*—similar in magnitude to that between *hb* and *kni*—prevents extended domain overlap in *D. melanogaster* (Figures 2F, 3B,D,F and Supplementary Figure S9; see also Supplementary Text S1). The balanced positive feedback loop implemented by these mutually inhibiting interactions maintains the *hb* boundary at a stable position (Figure 7). Taken together, this explains the counter-intuitive fact that weaker repression of *hb* by Kr (our model prediction) leads to an increased net interaction between the two genes in *M. abdita* [46] due to the larger overlap between domains in this species.

**Figure 3.**
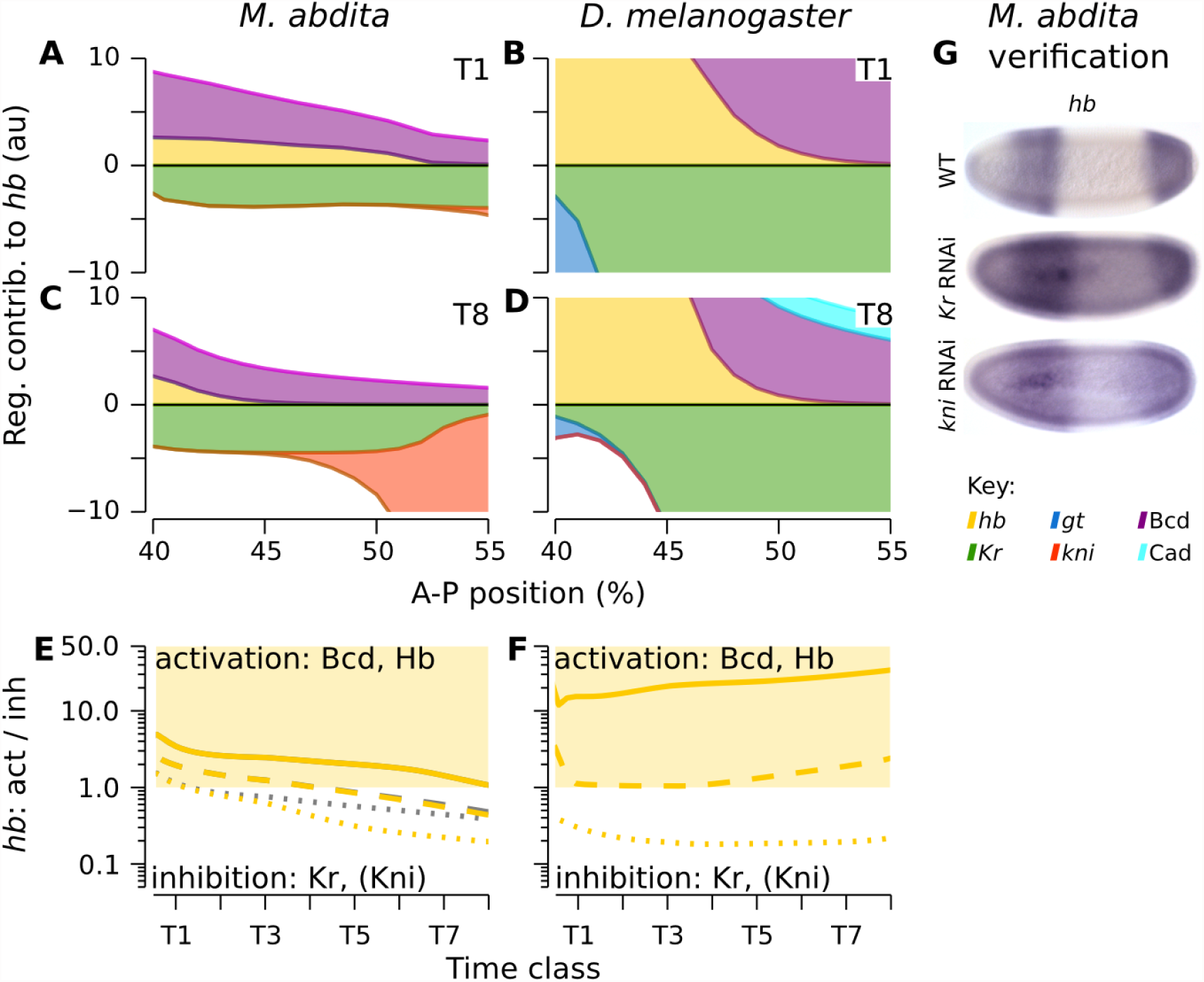
Graphical analysis of regulatory interactions involved in positioning the *hb* boundary. The left column of graphs shows *M. abdita*, the right column *D. melanogaster.* **(A–D)** Plots showing cumulative regulatory contributions of gap genes and external inputs to anterior *hb* in the region of its posterior boundary. Contributions are shown at time class T1 **(A, B)** and T8 **(C, D)**. Each coloured area corresponds to an individual regulatory term 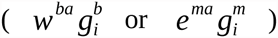 in Equation (3). Activating contributions are >0.0 and inhibiting contributions < 0.0. **(E, F)** Plots show ratios of activating vs. repressive regulatory input on *hb*, plotted over time for three equidistant nuclei at 40% (solid), 45% (dashed), and 50% (dotted) A–P position (grey lines exclude the additional repressor Kni, which is only active in the posterior-most nucleus at 50%). Light yellow areas indicate activation of *hb* (>1.0), white areas inhibition (< 1.0). Comparing curves in **E** vs. **F** reveals that Kr is sufficient to trigger *hb* down-regulation in *M. abdita*, but not in *D. melanogaster.* **(G)** Embryos of wild-type (WT) and RNAi-treated *M. abdita* embryos stained for *hb* mRNA at time class T5. Embryos are shown in lateral view: anterior is to the left, dorsal is up.

Our modeling results suggest that the position of the *hb* boundary depends on both Kr and Kni in *M. abdita*, while these factors act in a redundant manner in *D. melanogaster*. These predictions are confirmed by experimental evidence: *hb* expands posteriorly upon either *Kr* or *kni* knock-down in *M. abdita* (Figure 3G) [46]. In contrast, the absence of both factors is necessary to perturb the position of the *hb* boundary in *D. melanogaster* [63,71–74].

### Conserved mechanisms regulate the *Kr-kni* and *kni-gt* boundary interfaces

In contrast to the *hb* boundary described above, the borders of the abdominal *kni* domain and its overlapping companions—the posterior boundary of central *Kr* as well as the anterior boundary of posterior *gt*—exhibit anterior shifts that are conserved between *M. abdita* and *D. melanogaster* (Figure 2C,D) [46,57,66,75]. Accordingly, gene circuit analysis reveals that the regulatory mechanisms underlying these shifts are also largely conserved.

For the *Kr-kni* boundary interface, the anterior shift in border position is caused by a simple asymmetry in repressive interactions: *M. abdita* gene circuits show strong and increasing inhibition of *Kr* by Kni, while there is no repression of *kni* by Kr (Figure 2F, 4A, C; see also Supplementary Figure S10). This asymmetry is less pronounced in models for *D. melanogaster*, which employ additional *Kr* auto-repression and *kni* auto-activation to create the regulatory imbalance between the two genes (Figure 2F, 4B, D; Supplementary Figure S11). Such auto-regulatory contributions are unlikely to be biologically significant. We do not see these interactions in *D. melanogaster* gap gene circuits fitted to protein data [57,60,65] and there is no experimental evidence to support their existence [36]. In contrast, repressive imbalance between *Kr* and *kni* is strongly supported by experimental evidence in both species. While *Kr* expression expands posteriorly in *M. abdita* embryos treated with *kni* RNAi, no effect on *kni* is observed in *Kr* knock-down embryos (Figure 4E) [46]. Similarly, *Kr* has been reported to expand posteriorly in *kni* mutants of *D. melanogaster* [76–78] (although a recent quantitative study failed to detect this effect [74]) while *kni* expression is not affected in *Kr* mutants [74,79].

**Figure 4.**
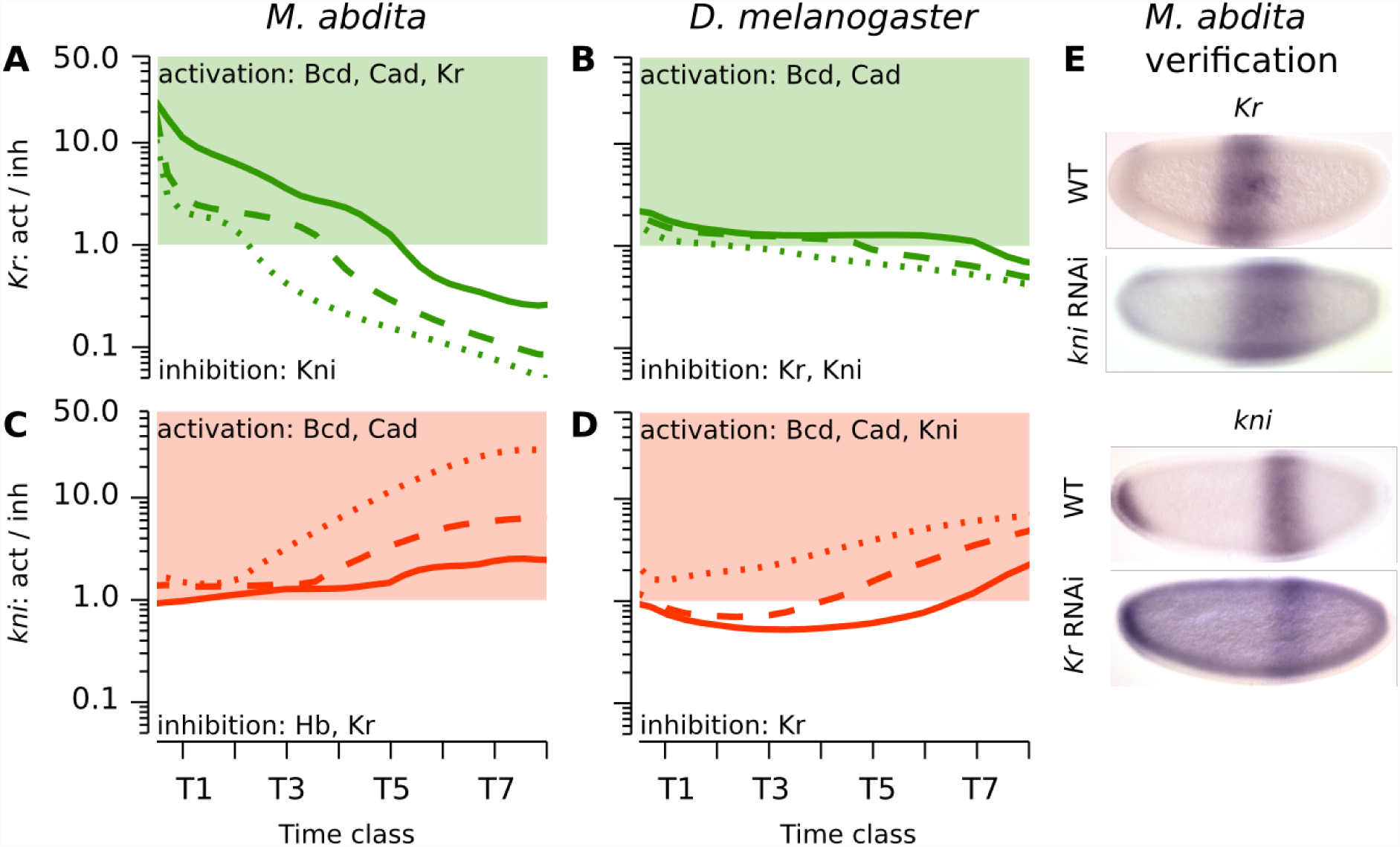
Graphical analysis of regulatory interactions involved in positioning the *Kr-kni* boundary interface. **(A–D)** Plots show ratios of activating vs. repressive regulatory input on *Kr* **(A, B)** and *kni* **(C, D)** over time in *M. abdita* **(A, C)** and *D. melanogaster* **(B, D)**. Lines indicate equidistant nuclei at 53% (solid), 55% (dashed), and 59% (dotted) A–P position **(A, C)**, and 54% (solid), 56% (dashed), and 58% (dotted) A–P position, respectively. Green/red coloured areas indicate activation of *Kr/kni* (>1.0), white areas indicate inhibition (< 1.0). Despite subtle differences in shift mechanism and dynamics, both *M. abdita* and *D. melanogaster* show increasing *Kr* repression and *kni* activation over time due to repressive asymmetry between the two genes. See main text for details. **(E)** Embryos of wild-type (WT) and RNAi-treated *M. abdita* embryos stained for *Kr* and *kni* mRNA at time class T3. Embryos are shown in lateral view: anterior is to the left, dorsal is up.

Regulation of the *kni-gt* boundary interface relies on an analogous repressive asymmetry and is also conserved between the two species, despite some differences in strength and timing of interactions: repression of *kni* by Gt is stronger than repression of *gt* by Kni in both species (Figure 2F, 5A–D; Supplementary Figures S12, S13). Experimental evidence supports these modeling predictions. The abdominal *kni* domain expands posteriorly in *M. abdita* embryos treated with *gt* RNAi, while posterior *gt* is not affected in *hb* RNAi knock-down embryos (Figure 5E) [46]. Similarly, the abdominal *kni* domain expands posteriorly in *gt* mutants of *D. melanogaster* [80]. In contrast, no effect on the anterior boundary of the posterior *gt* domain has been observed in *D. melanogaster kni* mutants [74,80–82].

**Figure 5.**
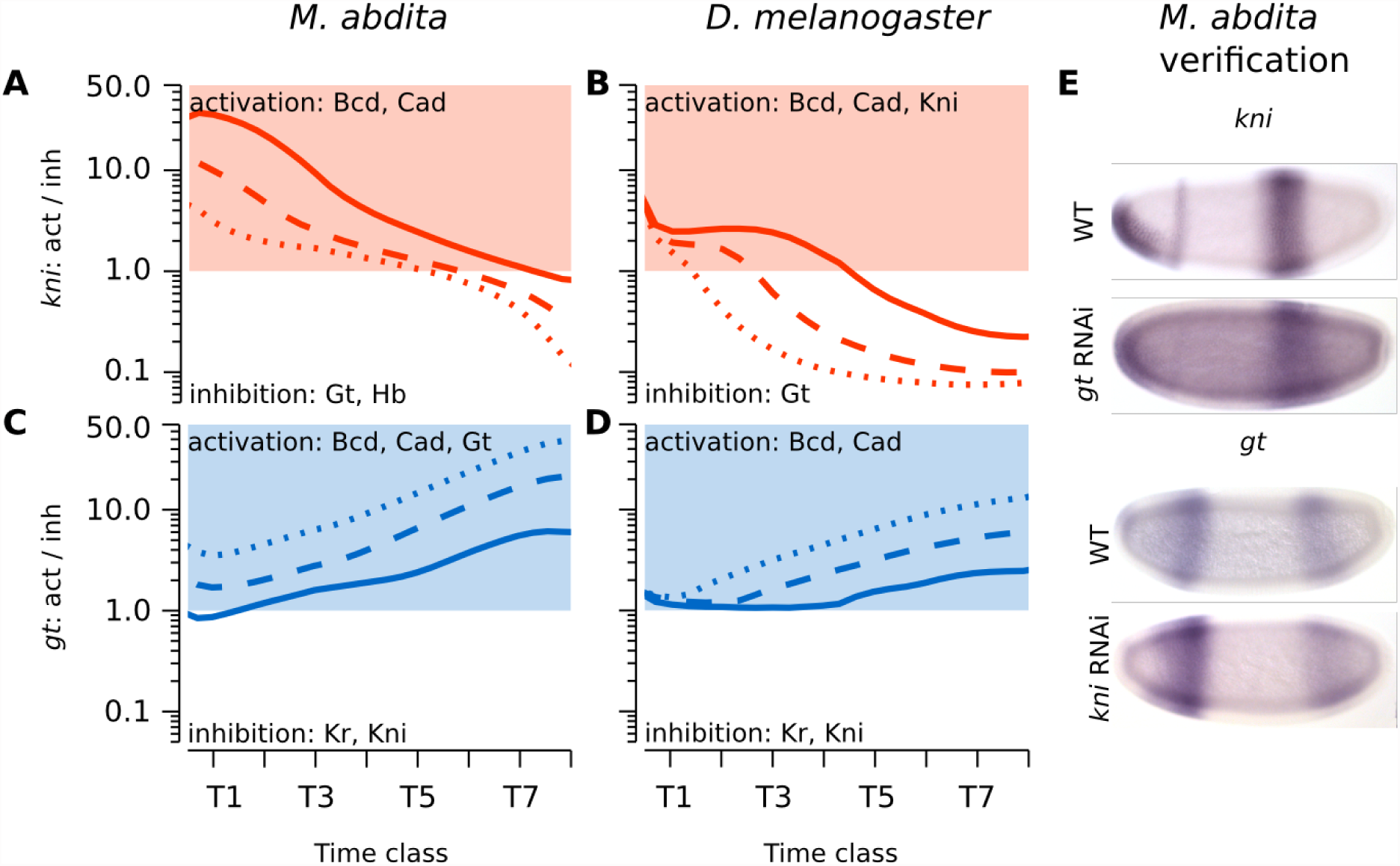
Graphical analysis of regulatory interactions involved in positioning the *kni-gt* boundary interface. (A–D) Plots show ratios of activating vs. repressive regulatory input on *kni* **(A, B)** and *gt* **(C, D)** over time in *M. abdita* **(A, C)** and *D. melanogaster* **(B, D)**. Lines indicate equidistant nuclei at 66% (solid), 68% (dashed), and 70% (dotted) A–P position **(A, C)**, and 65% (solid), 67% (dashed), and 69% (dotted) A–P position, respectively. Red/blue coloured areas indicate activation of *kni/gt* (>1.0), white areas indicate inhibition (< 1.0). Despite subtle differences in shift mechanism and dynamics, both *M. abdita* and *D. melanogaster* show increasing *kni* repression and *gt* activation over time due to repressive asymmetry between the two genes. See main text for details. **(E)** Embryos of wild-type (WT) and RNAi-treated *M. abdita* embryos stained for *kni* and *gt* mRNA at time class T5. Embryos are shown in lateral view: anterior is to the left, dorsal is up.

### Altered two-phase mechanism of posterior gap gene expression

Expression dynamics at the interface of the posterior *gt* and *hb* domains differs markedly between *M. abdita* and *D. melanogaster* (Figure 2C,D). In *D. melanogaster*, the posterior boundary of the posterior *gt* domain shifts at a constant rate over time. In contrast, this shift is delayed in *M. abdita*—due to the absence of maternal Cad [46,47,52]—until mid cleavage cycle 14A when it suddenly initiates and then proceeds much faster than in *D. melanogaster* (Figure 2D). Our models show that this behavior is governed through down-regulation of *gt* by Hb, whose posterior domain appears abruptly in *M. abdita* [46], while it accumulates gradually in *D. melanogaster* (Figure 6A–F). This dynamic discontinuity is caused by two distinct phases of *hb* regulation in *M. abdita* (Figure 6E; Supplementary Figure S14; see also summary in Figure 7 and Supplementary Text S1). In the first phase, activation by Gt (Figure 2F) boosts *hb* expression within an extended zone of domain overlap, until a threshold is reached which leads to a sudden increase in *hb* auto-activation. This initiates the second phase, in which *hb* acts to maintain its own expression, tilting the regulatory balance towards repression of *gt* by Hb. This “pull-and-trigger” temporal switch in activating contributions is not observed in *D. melanogaster*, where Gt represses *hb* and strong *hb* auto-activation is already active at earlier stages (Figure 6F, Supplementary Figure S15).

**Figure 6.**
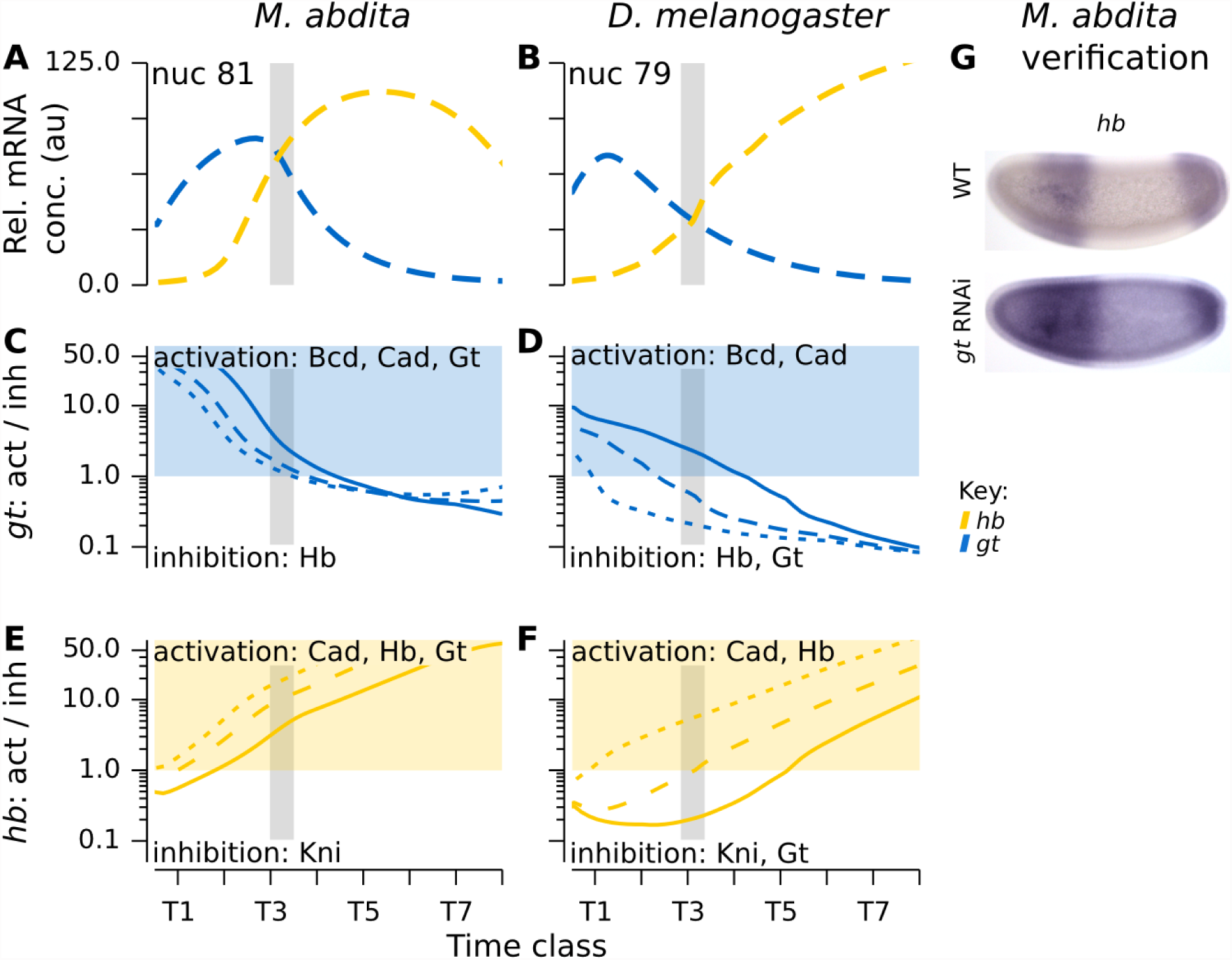
Graphical analysis of regulatory interactions involved in positioning the posterior *gt-hb* boundary interface. The left column of graphs shows *M. abdita*, the right column *D. melanogaster.* **(A, B)** Plots show relative mRNA concentrations of *gt* (blue) and *hb* (yellow) in nuclei at 81 **(A)** and 79% **(B)** A–P position. **(C–F)** Plots show ratios of activating vs. repressive regulatory input on *gt* **(C, D)** and *hb* **(E, F)** over time. Lines indicate equidistant nuclei at 79% (solid), 81% (dashed), and 83% (dotted) A–P position **(A, C)**, and 77% (solid), 79% (dashed), and 81% (dotted) A–P position, respectively. Blue/yellow coloured areas indicate activation of *gt/hb* (>1.0), white areas indicate inhibition (< 1.0). In *M. abdita*, down-regulation of *gt*, and concomitant up-regulation of *hb*, occur suddenly around mid cleavage cycle C14A (grey bar), while this process is much more gradual in *D. melanogaster.* **(E)** Embryos of wild-type (WT) and RNAi-treated *M. abdita* embryos stained for *hb* mRNA at time class T3. Embryos are shown in lateral view: anterior is to the left, dorsal is up.

These predictions are confirmed by experimental evidence: while the posterior *hb* domain is reduced in *M. abdita gt* knock-down embryos (Figure 3J) [46], no such effect can be seen in *gt* mutants of *D. melanogaster* [80,83]. In addition, our models clarify an ambiguous result from our experimental analysis [46], by establishing that the activation of *hb* by Gt is likely to be direct and functionally important.

### The role of domain overlaps and regulatory strength in cross-repression

Previous experimental evidence suggests that cross-repression between overlapping gap genes is stronger in *M. abdita*, since RNAi knock-downs show less ambiguous effects on posterior neighboring domains than the corresponding gap gene mutants in *D. melanogaster* [46]. In the case of *Kr* and *kni*, gene circuits confirm that this is caused by stronger asymmetry in the strength of regulatory interactions between these genes. In other cases, however, our models predict weaker gap-gap cross-repression. How can these apparently contradictory conclusions be reconciled?

The problem lies in the assumption that there is a direct and simple connection between sensitivity to RNAi knock-down and network interaction strength as represented by regulatory parameters. Our models, however, reveal a more intricate picture. For both Kr’s role in regulating *hb*, and the posterior *gt-hb* boundary interface, the relevant regulatory parameter values are smaller in *M. abdita* than in *D. melanogaster.* At first sight, this is puzzling. However, the problem is resolved if we consider that weaker repression allows for co-expression of gap genes across larger regions of the embryo. This is reflected in the expression data, which show that gap gene mRNA domains overlap much more extensively in *M. abdita* than in *D. melanogaster*, especially during the early stages of expression [46].

The proposed ratchet mechanism for placing the *hb* boundary, as well as the pull-and-trigger mechanism governing two-phase *gt-hb* dynamics in the posterior in *M. abdita*, both explicitly rely on extensive domain overlaps to function (see previous sections and summary in Figure 7). In contrast, the corresponding mechanisms in *D. melanogaster*, which are driven by positive feedback, prevent such overlap. In this way, our models reveal that sensitivity to genetic perturbations corresponds to the product of network interaction strength and the spatial extent to which regulators co-exist in the embryo.

**Figure 7.**
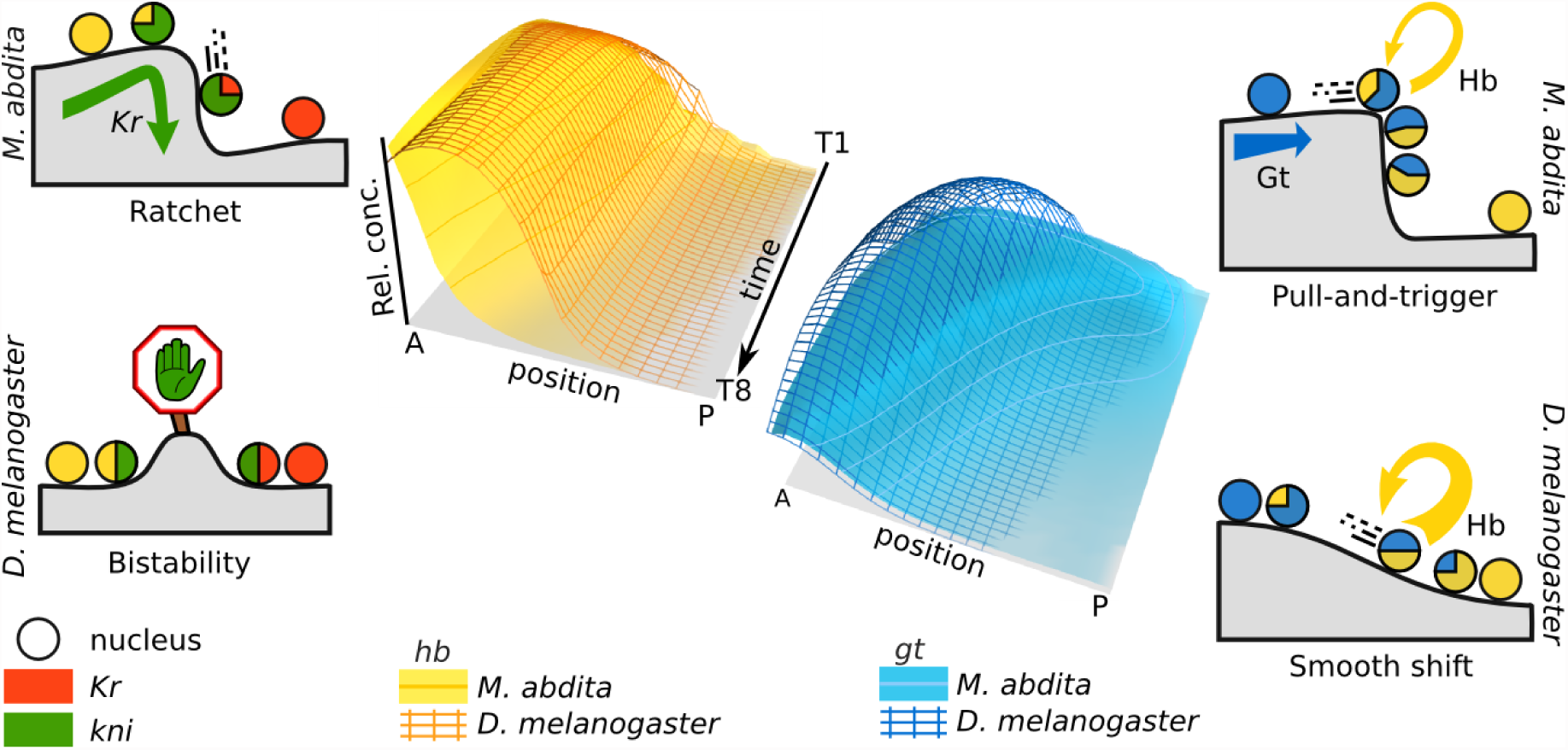
Divergent regulatory mechanisms for dynamic gap boundary placement. The gap gene networks of *M. abdita* and *D. melanogaster* exhibit quantitative differences in genetic interactions, which lead to qualitative differences in expression dynamics (shown as comparative 3D space-time plots). Cartoons illustrate the regulatory mechanisms underlying these differences: grey landscapes represent change in cell state; nuclei are shown as circles (color indicating the gap genes they express; colors as in Figs. 1–6). The posterior boundary of the anterior *hb* domain (left) is positioned by a “ratchet” mechanism in *M. abdita*: repression by Kr primes nuclei for a switch to strong repression by Kni resulting in an anterior shift of the *hb* boundary over time. In contrast, this boundary is set by a bistable switch mechanism based on mutual repression between *hb* and *Kr/kni* in *D. melanogaster*, resulting in a stationary boundary position. The posterior boundary of the posterior *gt* domain (right) is positioned through repression by Hb. In *M. abdita*, posterior *hb* is up-regulated in two phases, by a “pull-and-trigger” mechanism: initially *hb* is activated by Gt (the “pull”); later, auto-activation is “triggered” and becomes more dominant. In contrast, *hb* accumulates gradually in *D. melanogaster* due to stronger auto-activation at early stages. Yellow arrow indicates differences in the strength of *hb* auto-activation.

### Gap domain shifts are sufficient to account for compensatory evolution

We have previously shown that differences in gene expression dynamics—specifically, in the dynamics of gap domain shifts—enable the gap gene network to compensate for differences in upstream regulatory input from altered maternal gradients [46]. This leads to gap gene expression patterns that are almost equivalent in both *M. abdita* and *D. melanogaster* by the onset of gastrulation (Figure 2C, D). Our experimental work suggested that quantitative changes in gap-gap cross-repression are responsible for the observed differences in shift dynamics. *M. abdita* gap gene circuits allow us to go beyond such hypotheses in two important ways.

First, gap gene circuits provide explicit regulatory mechanisms for altered domain shifts. They give us causal rather than correlative explanations. Dynamic modeling allows us to explicitly track all simultaneous regulatory interactions across space and time. This cannot be achieved by experimental approaches alone.

Second, our models allow us to test whether the suggested changes in regulatory structure are necessary and sufficient to explain the observed changes in expression dynamics between *M. abdita* and *D. melanogaster*. Our analysis clearly demonstrates that this is indeed the case. They reveal that the most important contributions to compensatory regulation come from distinct mechanisms for the placement of the posterior boundary of anterior *hb* and the dynamic placement of the posterior *gt-hb* interface (Figure 7).

## Discussion and Conclusion

In this paper, we have provided a detailed comparative analysis—based on data-driven modeling and reverse engineering—of the regulatory mechanisms for compensatory evolution in the gap gene system of cyclorrhaphan flies. Our analysis provides causal-mechanistic explanations, in terms of dynamic regulatory mechanisms, for the observed differences in gap gene expression dynamics between *M. abdita* and *D. melanogaster.*

At first glance, the inter-species differences may appear subtle. However, the fact that we can capture and analyze such subtle changes demonstrates the sensitivity and accuracy of our quantitative approach. Moreover, small expression changes can be as important as large ones. In the case of the *hb* boundary, the change from stationary to moving boundary implies a qualitatively different dynamic regime for the underlying regulatory mechanism (Figure 7). Similarly, the dynamics of the posterior *gt-hb* interface involves two qualitatively different phases of dynamic regulation (Figure 7). We have shown that these altered mechanisms are sufficient to explain the observed compensatory dynamics. This kind of compensatory evolution leads to system drift [5,53–56]. It enables the gap gene networks of both species to produce equivalent patterning outputs despite differing maternal inputs [46,47].

In order for system drift to occur, there must be many different network “genotypes” (*i.e.* regulatory structures or GRNs) that produce the same “phenotype” (patterning outcome). Computer simulations of large ensembles of GRNs show that this is indeed the case [8,84,85]. Not only do such invariant sets of genotypes exist, but theoretical studies also show that most of the regulatory structures contained within them are connected by small mutational steps, forming what are called genotype networks [19,26,86,87]. A genotype network is a meta-network (a network of distinct GRNs producing the same phenotype) where each genotype is connected to another through the alteration of a single network interaction. Genotype networks provide the substrate for system drift: evolving regulatory networks can explore a genotype network, modifying and rewiring their structure as they go along, while maintaining a constant patterning output (see Box 1). Our models show that only slight changes to the strength of repressive interactions are sufficient to enable system drift. It is reasonable to assume that such changes can be achieved in relatively few mutational steps. In this way, our results indicate the presence of a genotype network underlying compensatory evolution of gap gene interactions.

System drift based on underlying genotype networks is not only an important mechanism for phenotypically neutral evolution, but is also an essential prerequisite for evolutionary innovation (and hence evolvability used in the narrow sense of the term, see Box 1) [19,26,33–35]. The reason for this is that different genotypes have different mutational neighborhoods. Only a subset of structural changes will maintain the output pattern and keep the system on its genotype network. Other mutations will lead to an altered (and potentially adaptive) novel phenotype. Network structures at different positions within a genotype network provide access to different phenotypes in their mutational neighborhood [19,26,30]. In this way, drift across a genotype network increases the diversity of accessible novel phenotypes, enabling the evolving system to explore new avenues of evolutionary change.

There is another way in which the regulatory structure of the gap gene network affects its evolvability. Our analysis reveals that some aspects of gap gene expression and regulation are more sensitive to parameter changes than others. The dynamics of domain shifts respond to subtle alterations in regulatory interaction strength. In contrast, the five main principles of gap gene regulation (shown in Figure 1B) are faithfully conserved among cyclorrhaphan flies despite considerable inter-species differences in the strength of regulatory interactions (Figure 2F) [46]. These results demonstrate how the regulatory structure of the gap gene network channels the direction of evolutionary change towards drift along the underlying genotype network. More generally, they show how random mutational changes lead to non-random changes in the patterning output of the system.

Our findings highlight the importance of dynamical systems theory for understanding regulatory evolution [10,11,14,30,64,88–90], in particular how a combination of differences in initial conditions (domain placement) and transient trajectories (expression dynamics) can explain compensatory changes in gene expression. More importantly, they show how subtle quantitative changes in the strength of regulatory interactions can give rise to qualitatively different regimes of expression dynamics (stationary vs. stable boundary; gradual vs. two-phase shift). The next step will be to understand how such transitions—and hence the evolutionary potential and evolvability of the system—can be explained by the geometry of the underlying configuration space of the models, that is to say by the arrangement of the system’s attractor states, their associated basins and their bifurcations [10,20,30,64,88–90].

Understanding such aspects of regulatory networks, in a quantitative and mechanistic manner, is essential if we are to move beyond the investigation of the role of individual genes towards elucidation of the dynamic principles governing regulatory evolution at the systems level [91]. The integrative approach we have presented here—based on data-driven modelling in non-model organisms [92]—provides a prototype for this kind of investigation.

## Materials and Methods

We infer the regulatory structure and dynamics of the gap gene network by means of gene circuits, dynamical network models that are fit to quantitative spatio-temporal expression data [57–61,65,66]. Here, we use the gene circuit approach with mRNA data—acquired and processed using efficient robust protocols and pipelines that work reliably in non-model species [93]. We have previously established that these kind of mRNA expression data are sufficient to successfully infer the gap gene network in *D. melanogaster* [66].

### Data Acquisition and Processing

#### Trunk Gap Genes

Gap gene circuits simulate expression and regulation of the four trunk gap genes *hb*, *Kr*, *gt*, and *kni*. Integrated mRNA expression data for these genes in *D. melanogaster* were published previously [66]. We constructed an equivalent integrated data set for *M. abdita* as follows.

Using a compound wide-field microscope, we took brightfield and DIC images of laterally oriented embryos stained for one or two gap genes using an enzymatic (colorimetric) *in situ* hybridisation protocol [66]. These images were then processed in three steps [93]: (1) We constructed binary whole-embryo masks by an edge-detection approach; using this mask, we rotated, cropped, and flipped the embryo images such that the A–P axis is horizontal, the anterior pole lies to the left, and dorsal is up; we then extracted raw gene expression intensities from a band along the lateral midline of the embryo covering 10% of the maximum dorso-ventral height. (2) We determined the position of gene expression domain boundaries by manually fitting clamped splines to the raw data. (3) Lastly, embryos were assigned to cleavage cycles C1–C14A based on the number of nuclei; cleavage cycle C14A was further subdivided into eight time classes (T1–T8, each about 7 min long) based on membrane morphology as described in [94]. Manual steps, such as spline fitting and time classification, are carried out by two researchers independently to detect and avoid bias. A detailed quantitative description and analysis of the resulting set of *M. abdita* gap gene expression boundaries is provided elsewhere [46,95].

We used the extracted domain boundaries to create an integrated spatio-temporal expression data set. We achieved this by computing median expression boundary positions for each gene and time class for which we have data [66]. During data processing, gap gene mRNA expression levels are normalised to the range [0.0, 1.0]. Because the gradual buildup and subsequent degradation of gap gene products is an important aspect of gap gene expression dynamics [62,67,75], we rescale these levels over space and time to create an expression data set that is comparable to previous mRNA data sets from *D. melanogaster* [66].

Embryo images, raw profiles, extracted boundaries, and integrated expression profiles for both fly species are available from the SuperFly database (http://superfly.crg.eu) [96].

#### External Inputs

Gap gene circuits require expression data for maternal co-ordinate genes *bcd* and *cad*, and the terminal gap genes *tll* and *hkb*, as external regulatory inputs. *D. melanogaster* data for these factors were described previously [66]. For *M. abdita*, see Extended Methods in S1 Supporting Information and Figure S1. In brief, the profile of *M. abdita* Cad protein is derived from immunostainings. Because we were unable to raise an antibody against *M. abdita* Bcd, we inferred its graded distribution through a simple model of protein diffusion from its localized mRNA source. We used mRNA data for *M. abdita tll* and *hkb* as we did for previous mRNA-based models for *D. melanogaster* [66].

### Gene Circuit Models

Gene circuits are mathematical models for simulating the regulatory dynamics of gene networks [57–61,65,66,97]. Gene circuits are hybrid models: continuous gene expression dynamics during interphase are complemented by discrete nuclear divisions between cleavage cycles.

Continuous gene regulatory dynamics are encoded by sets of ordinary differential equations (ODEs), each of which describes the change in concentration for a specific gene product *g* over time *t* in a particular nucleus *i* along the A–P axis (Figure 2B):

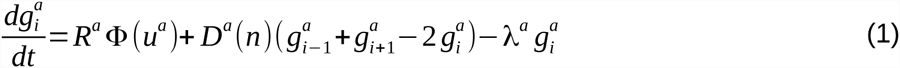

with *a*∈*G*, G = {*hb, Kr, gt, kni*}; regulated mRNA synthesis at maximum rate *R*; diffusion of gap gene products between neighboring nuclei (diffusion rate *D(n)* depends on nuclear density and hence the number of preceding mitoses *n*); and gene product degradation at rate *λ*. The saturating nature of gene regulation is captured by the sigmoid response function Φ(*u*^*a*^) :

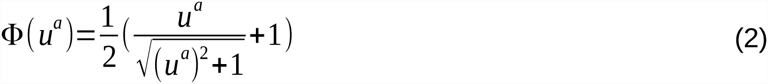

where

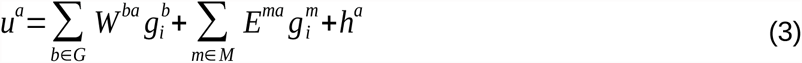

with the trunk gap genes G defined as above; the external inputs as M = {Bcd, Cad, Tll, Hkb}; and ubiquitous maternal activating or repressing factors represented by threshold parameter *h*. Interconnectivity matrices *W* and *E* define genetic interactions between the gap genes, and between external inputs and the gap genes, respectively.

Gene circuits cover the time from the initiation of gap gene expression to the onset of gastrulation: from C12 to the end of C14A (at *t = 98.667 minutes*) in *M. abdita*, from C13 to the end of C14A (at *t = 71.100 minutes*) in *D. melanogaster*. Mitotic division schedules are based on [94] for *M. abdita* and [66] for *D. melanogaster* (Supplementary Table S2). At each division, the number of nuclei, and hence the number of ODEs, doubles, while the distance between nuclei is halved. Nuclei are represented as a one-dimensional array along the A–P axis, covering the trunk region of the embryo (30–91% A–P position for *M. abdita*; 35–87% for *D. melanogaster*, 0% is at the anterior pole). Spatial ranges were chosen in accordance with earlier *D. melanogaster* models [65,66] to represent an equivalent set of a gap domains in each species.

### Model Fitting

We fit gene circuit models to quantitative expression data from both *M. abdita* and *D. melanogaster* as described previously [66]. In brief, the values of gene circuit parameters are estimated by means of a robust global optimisation algorithm called parallel Lam Simulated Annealing (pLSA) [98]. pLSA iteratively approximates the minimum of a cost function which represents the difference between model output and expression profiles in the data (Figure 2B). We have previously established that a Weighted Least Squares (WLS) cost function—with artificial weights that are inversely proportional to expression levels, thus strongly penalizing ectopic expression—is effective at fitting gene circuits to mRNA gap gene expression data in *D. melanogaster* [65,66]. Further details on the optimization procedure can be found in Supporting Information.

Model fitting was performed on the Mare Nostrum supercomputer at the Barcelona Supercomputing Centre (http://www.bsc.es). The average duration of a pLSA run on 64 cores is approximately 45 min. For *M. abdita*, we performed series of global optimization runs—comprising 1650 independent model fits in total—that cover a number of different scenarios: estimated Bcd gradients with different scales, circuits with or without diffusion or auto-regulation, and circuits fitted while not fixing threshold parameters *h* (see Supplementary Text S1, Table S3). We also obtained a reference set of 225 *D. melanogaster* gap gene circuits. These models differ slightly from those published previously [66] since they use an approximation for the Bcd gradient equivalent to that used in gene circuits for *abdita*.

### Selection of Gene Circuits for Analysis

All 225 *D. melanogater* circuits, and the best-fitting scenario for *M. abdita* (400 circuits; see Supplementary Methods), were chosen for further analysis. These circuits were then subjected to the following quality tests. Numerically unstable circuits were discarded, as were all fits with an root-mean-square score larger than 30.0 [66]. The remaining gene circuits were visually inspected for defects in gene expression profiles (see Supplementary Methods, for details). This selection process resulted in 20 solutions in each species. Their expression dynamics are shown in Supplementary Fig. S5.

### Computational Tools

Image processing and extraction/measurement of expression domain boundary positions was performed with the Java application FlyGUI (https://subversion.assembla.com/svn/flygui) [93]. Our gene expression data sets are available from the SuperFly website (http://www.superfly.crg.es) [96], and from Figshare [95]. Simulation and optimisation code is implemented in C, using MPI for parallelization, SUNDIALS (http://computation.llnl.gov/casc/sundials) for numerical solvers [99], and the GNU Scientific Library (GSL, http://www.gnu.org/software/gsl) for data interpolation (https://subversion.assembla.com/svn/flysa).

## Acknowledgements

We thank Urs Schmidt-Ott and Steffen Lemke for crucial help and support to get us off the ground with *M. abdita*; Brenda Gavilán and Núria Bosch Guiteras for help with maintaining the *M. abdita* culture; and Damjan Cicin-Sain, Toni Hermoso Pulido, and the CRG Bioinformatics Core for computational support. John Reinitz, James Sharpe, Ben Lehner, Erik Clark, Hilde Janssens, Berta Verd, and Simone Reber provided useful feedback on the manuscript. Adam Wilkins provided essential editing and intellectual arguments to improve the presentation of our argument. We thankfully acknowledge the computer resources, technical expertise and assistance provided by the Barcelona Supercomputing Center—Centro Nacional de Supercomputación. This work was funded by the MEC-EMBL agreement for the EMBL/CRG Research Unit in Systems Biology, European Commission grant FP7-KBBE-2011-5/289434 (BioPreDyn), by Grant 153 (MOPDEV) of the ERANet: ComplexityNET program, AGAUR SGR Grant 406, as well as grants BFU2009-10184 and BFU2012-33775 from MINECO. The Centre for Genomic Regulation (CRG) acknowledges support from MINECO, ‘Centro de Excelencia Severo Ochoa 2013-2017’, SEV-2012-0208.

## Author Contributions

AC fitted and analyzed the models, and contributed to data processing/quantification; KRW produced and processed *M. abdita* quantitative data sets and performed RNAi knock-downs, with substantial contributions from EJG; JJ conceived the study and supervised experimental/computational work; AC and JJ wrote the paper, with contributions from KRW and EJG.

## Supporting Information

S1 Text. Extended methods and results.

S1 Figure. Estimated protein expression patterns of maternal co-ordinate genes *bcd* (purple) and *cad* (cyan).

S2 Figure. Genetic interconnectivity matrices for scenarios with different *M. abdita* Bcd gradients. S3 Figure. Genetic interconnectivity matrices for *D. melanogaster* gene circuits fit to mRNA or protein data.

S4 Figure. Common gene expression defects in *M. abdita* gene circuits. S5 Figure. Gene circuit fits to data in *M. abdita* and *D. melanogaster*.

S6 Figure. Comparison of interaction strengths for all regulatory weights across species.

S7 Figure. Comparison of maximum production rates, diffusion parameters, and gene product half lives between species.

S8 Figure. Graphical analysis of the *hb-Kr* interface in *M. abdita*.

S9 Figure. Graphical analysis of the *hb-Kr* interface in *D. melanogaster*. S10 Figure. Graphical analysis of the *Kr-kni* interface in *M. abdita*.

S11 Figure. Graphical analysis of the *Kr-kni* interface in *D. melanogaster*. S12 Figure. Graphical analysis of the *kni-gt* interface in *M. abdita*.

S13 Figure. Graphical analysis of the *kni-gt* interface in *D. melanogaster*. S14 Figure. Graphical analysis of the *gt-hb* interface in *M. abdita*.

S15 Figure. Graphical analysis of the *gt-hb* interface in *D. melanogaster*. S1 Table. Extent of the *bcd* mRNA domain.

S2 Table. Mitotic division schedules and time classes for *M. abdita* and *D. melanogaster*. S3 Table. Selection of gene circuits for analysis.

S4 Table. Presence of patterning mechanisms in *M. abdita* scenarios and *D. melanogaster* reference gene circuits.

## References

1. Jaeger J, Sharpe J. On the concept of mechanism in development. In: Minelli A, Pradeu T, editors. Towards a Theory of Development. Oxford: Oxford University Press; 2014. p. 56.

2. Smith JM, Burian R, Kauffman S, Alberch P, Campbell J, Goodwin B, et al. Developmental Constraints and Evolution: A Perspective from the Mountain Lake Conference on Development and Evolution. Q Rev Biol. 1985;60: 265. doi:10.1086/414425

3. Wilkins AS. Between “design” and “bricolage”: genetic networks, levels of selection, and adaptive evolution. Proc Natl Acad Sci U S A. 2007;104 Suppl 1: 8590–8596. doi:10.1073/pnas.0701044104

4. Salazar-Ciudad I, Marín-Riera M. Adaptive dynamics under development-based genotype-phenotype maps. Nature. 2013;497: 361–4. doi:10.1038/nature12142

5. Pavlicev M, Wagner GP. A model of developmental evolution: Selection, pleiotropy and compensation. Trends Ecol Evol. 2012;27: 316–322. doi:10.1016/j.tree.2012.01.016

6. Arthur W. The emerging conceptual framework of evolutionary developmental biology. Nature. 2002;415: 757–764. doi:10.1038/415757a

7. Counce SJ. The Strategy of the Genes. The Yale journal of biology and medicine. London: George Allen & Unwin; 1958.

8. Kauffman S a. Metabolic stability and epigenesis in randomly constructed genetic nets. J Theor Biol. 1969;22: 437–467. doi:10.1016/0022-5193(69)90015-0

9. J. R, J. R. Regulation Cells: Science. 1969; 349–357.

10. Thom R. Structural stability and morphogenesis. Pattern Recognition. Perseus Books Group; 1976. doi:10.1016/0031-3203(76)90030-3

11. Group DB, Sciences B. and Evolution. Dev Biol. 1982;97: 43–55. doi:10.1016/j.semcancer.2013.09.001.B-cell

12. Alberch P. From genes to phenotype: dynamical systems and evolvability. Genetica. 1991;84: 5–11. doi:10.1007/BF00123979

13. Kauffman SA. The origins of order; self organization and selection in evolution. International Journal of Biochemistry. Oxford University Press; 1994. doi:10.1016/0020-711X(94)90119-8

14. Webster G, Goodwin BC. Form and transformation: generative and relational principles in biology. Cambridge: Cambridge University Press; 1996.

15. Wilkins AS. Recasting developmental evolution in terms of genetic pathway and network evolution … and the implications for comparative biology. Brain Research Bulletin. 2005. pp. 495–509. doi:10.1016/j.brainresbull.2005.04.001

16. Davidson EH, Erwin DH. Gene regulatory networks and the evolution of animal body plans. Science. 2006;311: 796–800. doi:10.1126/science.1113832

17. Wilkins AS. Genetic networks as transmitting and amplifying devices for natural genetic tinkering. Novartis Found Symp. 2007;284: 71–86; discussion 86–89, 110–115.

18. Wilke CO. Robustness and Evolvability in Living Systems. BioScience. 2006. doi:10.1641/0006-3568(2006)56[695:RAEILS]2.0.CO;2

19. Wagner A. Robustness and evolvability: a paradox resolved. Proc Biol Sci. 2008;275: 91–100. doi:10.1098/rspb.2007.1137

20. Crombach A, Hogeweg P. Evolution of evolvability in gene regulatory networks. PLOS Comput Biol. 2008;4: e1000112. doi:10.1371/journal.pcbi.1000112

21. Stern DL, Orgogozo V. The loci of evolution: How predictable is genetic evolution? Evolution. 2008. pp. 2155–2177. doi:10.1111/j.1558-5646.2008.00450.x

22. Stern DL, Orgogozo V. Is genetic evolution predictable? Science. 2009;323: 746–751. doi:10.1126/science.1158997

23. Erwin DH, Davidson EH. The evolution of hierarchical gene regulatory networks. Nat Rev Genet. 2009;10: 141–148. doi:10.1038/nrg2499

24. Draghi J a, Parsons TL, Wagner GP, Plotkin JB. Mutational robustness can facilitate adaptation. Nature. 2010;463: 353–355. doi:10.1038/nature08694

25. Cotterell J, Sharpe J. An atlas of gene regulatory networks reveals multiple three-gene mechanisms for interpreting morphogen gradients. Mol Syst Biol. 2010;6: 425. doi:10.1038/msb.2010.74

26. Wagner A. The Origins of Evolutionary Innovations: A Theory of Transformative Change in Living Systems. Oxford University Press; 2011.

27. Peter I, Davidson E. Evolution of Gene Regulatory Networks that Control Embryonic Development of the Body Plan. Cell. 2011;144: 970–985. doi:10.1016/j.cell.2011.02.017. Evolution

28. Hoyos E, Kim K, Milloz J, Barkoulas M, Pénigault JB, Munro E, et al. Quantitative variation in autocrine signaling and pathway crosstalk in the *Caenorhabditis* vulval network. Curr Biol. 2011;21: 527–538. doi:10.1016/j.cub.2011.02.040

29. Wagner GP. Homology, Genes, and Evolutionary Innovation. Princeton University Press; 2014.

30. Jiménez A, Munteanu A, Sharpe J. Dynamics of gene circuits shapes evolvability. Proc Natl Acad Sci. 2015; 201411065. doi:10.1073/pnas.1411065112

31. Sorrells TR, Johnson AD. Making Sense of Transcription Networks. Cell. Elsevier Inc.; 2015;161: 714–723. doi:10.1016/j.cell.2015.04.014

32. Peter I, Davidson EH. Genomic Control Process. Oxford: Academic Press; 2015.

33. Wagner GP, Altenberg L. Perspective: Complex adaptations and the evolution of evolvability. Evolution (N Y). 1996;50: 967–976. doi:10.2307/2410639

34. Hendrikse JL, Parsons TE, Hallgrímsson B. Evolvability as the proper focus of evolutionary developmental biology. Evol Dev. 2007;9: 393–401. doi:10.1111/j.1525-142X.2007.00176.x

35. Pigliucci M. Is evolvability evolvable? Nat Rev Genet. 2008;9: 75–82. doi:10.1038/nrg2278

36. Jaeger J. The gap gene network. Cell Mol Life Sci. Springer; 2011;68: 243–274. doi:10.1007/s00018-010-0536-y

37. Sommer R, Tautz D. Segmentation gene expression in the housefly *Musca domestica*. Development. 1991;113: 419–430.

38. Rohr KB, Tautz D, Sander K. Segmentation gene expression in the mothmidge *Clogmia albipunctata* (Diptera, Psychodidae) and other primitive dipterans. Dev Genes Evol. 1999;209: 145–154. doi:10.1007/s004270050238

39. Stauber M, Taubert H, Schmidt-Ott U. Function of *bicoid* and *hunchback* homologs in the basal cyclorrhaphan fly *Megaselia* (Phoridae). Proc Natl Acad Sci U S A. 2000;97: 10844–10849. doi:10.1073/pnas.190095397

40. McGregor a. P, Shaw PJ, Dover G a. Sequence and expression of the *hunchback* gene in *Lucilia sericata*: A comparison with other Dipterans. Dev Genes Evol. 2001;211: 315–318. doi:10.1007/s004270100148

41. Goltsev Y, Hsiong W, Lanzaro G, Levine M. Different combinations of gap repressors for common stripes in *Anopheles* and *Drosophila* embryos. Dev Biol. 2004;275: 435–446. doi:10.1016/j.ydbio.2004.08.021

42. Hare EE, Peterson BK, Iyer VN, Meier R, Eisen MB. Sepsid *even-skipped* enhancers are functionally conserved in *Drosophila* despite lack of sequence conservation. PLoS Genet. 2008;4. doi:10.1371/journal.pgen.1000106

43. Lemke S, Schmidt-Ott U. Evidence for a composite anterior determinant in the hover fly *Episyrphus balteatus* (Syrphidae), a cyclorrhaphan fly with an anterodorsal serosa anlage. Development. 2009;136: 117–127. doi:10.1242/dev.033530

44. Lemke S, Busch SE, Antonopoulos D a, Meyer F, Domanus MH, Schmidt-Ott U. Maternal activation of gap genes in the hover fly *Episyrphus*. Development. 2010;137: 1709–1719. doi:10.1242/dev.055558

45. Crombach A, García-Solache M a., Jaeger J. Evolution of early development in dipterans: Reverse-engineering the gap gene network in the moth midge *Clogmia albipunctata* (Psychodidae). BioSystems. 2014;123: 74–85. doi:10.1016/j.biosystems.2014.06.003

46. Wotton KR, Jiménez-Guri E, Crombach A, Janssens H, Alcaine-Colet A, Lemke S, et al. Quantitative system drift compensates for altered maternal inputs to the gap gene network of the scuttle fly *Megaselia abdita*. Elife. 2015;4. doi:10.7554/eLife.04785

47. Wotton KR, Jiménez-Guri E, Jaeger J. Maternal Co-ordinate Gene Regulation and Axis Polarity in the Scuttle Fly *Megaselia abdita*. PLoS Genet. 2015;11: e1005042. doi:10.1371/journal.pgen.1005042

48. García-Solache M, Jaeger J, Akam M. A systematic analysis of the gap gene system in the moth midge *Clogmia albipunctata*. Dev Biol. Elsevier Inc.; 2010;344: 306–318. doi:10.1016/j.ydbio.2010.04.019

49. Klomp J, Athy D, Kwan CW, Bloch NI, Sandmann T, Lemke S, et al. A cysteine-clamp gene drives embryo polarity in the midge *Chironomus*. Science. 2015;348: 1040–1042. doi:10.1126/science.aaa7105

50. Rafiqi AM, Lemke S, Schmidt-Ott U. The scuttle fly *Megaselia abdita* (Phoridae): A link between *Drosophila* and mosquito development. Cold Spring Harb Protoc. 2011;6: pdb.emo143. doi:10.1101/pdb.emo143

51. Stauber M, Jäckle H, Schmidt-Ott U. The anterior determinant *bicoid* of *Drosophila* is a derived Hox class 3 gene. Proc Natl Acad Sci U S A. 1999;96: 3786–3789. doi:10.1073/pnas.96.7.3786

52. Stauber M, Lemke S, Schmidt-Ott U. Expression and regulation of *caudal* in the lower cyclorrhaphan fly *Megaselia*. Dev Genes Evol. 2008;218: 81–87. doi:10.1007/s00427-008-0204-5

53. True JR, Haag ES. Developmental system drift and flexibility in evolutionary trajectories. Evol Dev. 2001;3: 109–119. doi:10.1046/j.1525-142X.2001.003002109.x

54. Haag ES. Compensatory vs. pseudocompensatory evolution in molecular and developmental interactions. Genetica. 2007;129: 45–55. doi:10.1007/s10709-006-0032-3

55. Weiss KM, Fullerton SM. Phenogenetic drift and the evolution of genotype-phenotype relationships. Theor Popul Biol. 2000;57: 187–195. doi:10.1006/tpbi.2000.1460

56. Weiss KM. The phenogenetic logic of life. Nat Rev Genet. 2005;6: 36–45. doi:10.1038/nrg1502

57. Jaeger J, Surkova S, Blagov M, Janssens H, Kosman D, Kozlov KN, et al. Dynamic control of positional information in the early *Drosophila* embryo. Nature. 2004;430: 368–371. doi:10.1038/nature02678

58. Reinitz J, Sharp DH. Mechanism of *eve* stripe formation. Mech Dev. 1995;49: 133–158. doi:10.1016/0925-4773(94)00310-J

59. Reinitz J, Mjolsness E, Sharp DH. Model for cooperative control of positional information in *Drosophila* by *bicoid* and maternal *hunchback*. J Exp Zool. 1995;271: 47–56. doi:10.1002/jez.1402710106

60. Jaeger J, Blagov M, Kosman D, Kozlov KN, Manu, Myasnikova E, et al. Dynamical analysis of regulatory interactions in the gap gene system of *Drosophila melanogaster*. Genetics. 2004;167: 1721–1737. doi:10.1534/genetics.104.027334

61. Perkins TJ, Jaeger J, Reinitz J, Glass L. Reverse engineering the gap gene network of *Drosophila melanogaster*. PLoS Comput Biol. 2006;2: e51. doi: 10.1371/journal.pcbi.0020051

62. Jaeger J, Sharp DH, Reinitz J. Known maternal gradients are not sufficient for the establishment of gap domains in *Drosophila melanogaster*. Mech Dev. 2007;124: 108–128. doi:10.1016/j.mod.2006.11.001

63. Manu, Surkova S, Spirov A V., Gursky I V., Janssens H, Kim AR, et al. Canalization of gene expression in the *Drosophila* blastoderm by gap gene cross regulation. PLoS Biol. 2009;7: 0591–0603. doi:10.1371/journal.pbio.1000049

64. Manu, Surkova S, Spirov A V., Gursky V V., Janssens H, Kim AR, et al. Canalization of gene expression and domain shifts in the *Drosophila* blastoderm by dynamical attractors. PLoS Comput Biol. 2009;5: e1000303. doi:10.1371/journal.pcbi.1000303

65. Ashyraliyev M, Siggens K, Janssens H, Blom J, Akam M, Jaeger J. Gene circuit analysis of the terminal gap gene *huckebein*. PLoS Comput Biol. 2009;5: e1000548. doi:10.1371/journal.pcbi.1000548

66. Crombach A, Wotton KR, Cicin-Sain D, Ashyraliyev M, Jaeger J. Efficient reverse-engineering of a developmental gene regulatory network. PLoS Comput Biol. 2012;8: e1002589. doi:10.1371/journal.pcbi.1002589

67. Becker K, Balsa-Canto E, Cicin-Sain D, Hoermann A, Janssens H, Banga JR, et al. Reverse-Engineering Post-Transcriptional Regulation of Gap Genes in *Drosophila melanogaster*. PLoS Comput Biol. 2013;9: e1003281. doi:10.1371/journal.pcbi.1003281

68. Gregor T, Tank DW, Wieschaus EF, Bialek W. Probing the Limits to Positional Information. Cell. 2007;130: 153–164. doi:10.1016/j.cell.2007.05.025

69. Liu F, Morrison AH, Gregor T. Dynamic interpretation of maternal inputs by the *Drosophila* segmentation gene network. Proc Natl Acad Sci U S A. 2013;110: 6724–9. doi:10.1073/pnas.1220912110

70. Liu J, Ma J. Uncovering a Dynamic Feature of the Transcriptional Regulatory Network for Anterior-Posterior Patterning in the *Drosophila* Embryo. PLoS One. 2013;8: e62641. doi:10.1371/journal.pone.0062641

71. Houchmandzadeh B, Wieschaus E, Leibler S. Establishment of developmental precision and proportions in the early *Drosophila* embryo. Nature. 2002;415: 798–802. doi:10.1038/415798a

72. Clyde DE, Corado MSG, Wu X, Paré A, Papatsenko D, Small S. A self-organizing system of repressor gradients establishes segmental complexity in *Drosophila*. Nature. 2003;426: 849–853. doi:10.1038/nature02189

73. Perry MW, Bothma JP, Luu RD, Levine M. Precision of *hunchback* expression in the *Drosophila* embryo. Curr Biol. 2012;22: 2247–2252. doi:10.1016/j.cub.2012.09.051

74. Surkova S, Golubkova E, Manu, Panok L, Mamon L, Reinitz J, et al. Quantitative dynamics and increased variability of segmentation gene expression in the *Drosophila Krüppel* and *knirps* mutants. Dev Biol. Elsevier; 2013;376: 99–112. doi:10.1016/j.ydbio.2013.01.008

75. Surkova S, Kosman D, Kozlov K, Manu, Myasnikova E, Samsonova A a., et al. Characterization of the *Drosophila* segment determination morphome. Dev Biol. 2008;313: 844–862. doi:10.1016/j.ydbio.2007.10.037

76. Jäckle H, Tautz D, Schuh R, Seifert E, Lehmann R. Cross-regulatory interactions among the gap genes of *Drosophila*. Nature. 1986;324: 668–670. doi:10.1038/324668a0

77. Gaul U, Seifert E, Schuh R, Jäckle H. Analysis of Krüppel protein distribution during early *Drosophila* development reveals posttranscriptional regulation. Cell. 1987;50: 639–647. doi:10.1016/0092-8674(87)90037-7

78. Harding K, Levine M. Gap genes define the limits of *antennapedia* and *bithorax* gene expression during early development in *Drosophila*. EMBO J. 1988;7: 205–214.

79. Capovilla M, Eldon ED, Pirrotta V. The *giant* gene of *Drosophila* encodes a b-ZIP DNA-binding protein that regulates the expression of other segmentation gap genes. Development. 1992;114: 99–112.

80. Eldon ED, Pirrotta V. Interactions of the *Drosophila* gap gene *giant* with maternal and zygotic pattern-forming genes. Development. 1991;111: 367–378.

81. Kraut R, Levine M. Spatial regulation of the gap gene *giant* during *Drosophila* development. Development. 1991;111: 601–609.

82. Mohler J, Eldon ED, Pirrotta V. A novel spatial transcription pattern associated with the segmentation gene, *giant*, of *Drosophila*. EMBO J. 1989;8: 1539–1548.

83. Strunk B, Struffi P, Wright K, Pabst B, Thomas J, Qin L, et al. Role of CtBP in transcriptional repression by the *Drosophila giant* protein. Dev Biol. 2001;239: 229–240. doi:10.1006/dbio.2001.0454

84. Borenstein E, Krakauer DC. An end to endless forms: Epistasis, phenotype distribution bias, and nonuniform evolution. PLoS Comput Biol. 2008;4. doi:10.1371/journal.pcbi.1000202

85. Munteanu A, Solé R V. Neutrality and robustness in evo-devo: Emergence of lateral inhibition. PLoS Comput Biol. 2008;4. doi:10.1371/journal.pcbi.1000226

86. Ciliberti S, Martin OC, Wagner A. Robustness can evolve gradually in complex regulatory gene networks with varying topology. PLoS Comput Biol. 2007;3: 0164–0173. doi:10.1371/journal.pcbi.0030015

87. Ciliberti S, Martin OC, Wagner a. Innovation and robustness in complex regulatory gene networks. Proc Natl Acad Sci U S A. 2007;104: 13591–13596. doi:10.1073/pnas.0705396104

88. François P, Siggia ED. Phenotypic models of evolution and development: Geometry as destiny. Current Opinion in Genetics and Development. 2012. pp. 627–633. doi:10.1016/j.gde.2012.09.001

89. Jaeger J, Irons D, Monk N. The Inheritance of Process: A Dynamical Systems Approach. J Exp Zool Part B. 2012;318: 591–612. doi:10.1002/jez.b.22468

90. Jaeger J, Monk N. Bioattractors: dynamical systems theory and the evolution of regulatory processes. J Physiol. 2014;592: 2267–81. doi:10.1113/jphysiol.2014.272385

91. Jaeger J, Laubichler M, Callebaut W. The Comet Cometh: Evolving Developmental Systems. Biol Theory. 2015;10: 36–49. doi:10.1007/s13752-015-0203-5

92. Jaeger J, Crombach A. Life’s Attractors. Evolutionary systems biology. Soyer, O editor. Springer New York; 2012. pp. 93–119.

93. Crombach A, Cicin-Sain D, Wotton KR, Jaeger J. Medium-Throughput Processing of Whole Mount *In Situ* Hybridisation Experiments into Gene Expression Domains. PLoS One. 2012;7: e46658. doi:10.1371/journal.pone.0046658

94. Wotton KR, Jiménez-Guri E, García Matheu B, Jaeger J. A staging scheme for the development of the scuttle fly *Megaselia abdita.* PLoS One. 2014;9: e84421. doi:10.1371/journal.pone.0084421

95. Wotton KR, Jiménez-Guri E, Crombach A, Cicin-Sain D, Jaeger J. High-resolution gene expression data from blastoderm embryos of the scuttle fly *Megaselia abdita*. Sci Data. 2015;2: 150005. doi:10.1038/sdata.2015.5

96. Cicin-Sain D, Pulido a. H, Crombach a., Wotton KR, Jimenez-Guri E, Taly J-F, et al. SuperFly: a comparative database for quantified spatio-temporal gene expression patterns in early dipteran embryos. Nucleic Acids Res. 2014;43: D751–D755. doi:10.1093/nar/gku1142

97. Mjolsness E, Sharp DH, Reinitz J. A connectionist model of development. Journal of Theoretical Biology. 1991. pp. 429–453. doi:10.1016/S0022-5193(05)80391-1

98. Chu K-W, Deng Y, Reinitz J. Parallel Simulated Annealing by Mixing of States. J Comput Phys. 1999;148: 646–662. doi:10.1006/jcph.1998.6134

99. Hindmarsh AC, Brown PN, Grant KE, Lee SL, Serban R, Shumaker DE, et al. SUNDIALS: Suite of Nonlinear and Differential/Algebraic Equation Solvers. ACM Trans Math Softw. 2005;31: 363–396. doi:10.1145/1089014.1089020

100. Machamer P, Darden L, Craver CF. Thinking about Mechanisms. Philosophy of Science. 2000. p. 1. doi:10.1086/392759

